# Optimization of Injection-Molded Thermoplastic Microfluidic Chip Design with Numerical Modeling and Two-Photon Polymerization 3D Printing

**DOI:** 10.1101/2025.07.23.666150

**Authors:** Calum Mallorie, Barna Schneeweis, Maximilian Pitzek, Christoph Holzner, Claudia Lidl, Sinead de Cleir, Nemanja Milanovic, Domenico Foglia, Timm Krüger, Thomas R. Carey

**Affiliations:** School of Engineering, University of Edinburgh, Edinburgh, Scotland, UK; Stratec Consumables GmbH, Salzburg, AT; Cytovale Inc., South San Francisco, CA, US; NanoVoxel GmbH, Vienna, AT

**Author notes:** Correspondence and requests for materials should be addressed to Thomas Carey.

## Abstract

Precise microfluidic geometries are critical for particle dynamics studies but challenging to fabricate repeatably and rigidly. This study aims to experimentally validate computational fluid dynamics predictions of particle trajectories in cross-slot junctions, overcoming previous manufacturing limitations. We developed a novel fabrication process combining precisely bonded injection-molded chips with integrated two-photon polymerization 3D-printed junction geometries. Using this approach, experimental measurements of microsphere trajectories via the Cytovale system successfully validated predictions, confirming the influence of stenosis and outlet widths. This work establishes a robust methodology for optimizing microfluidic designs by synergizing advanced manufacturing and simulation, enabling precise experimental investigations.

## 1 Introduction

Cross-slot cytometry is a powerful tool for measuring particle properties with applications in diagnostics,^1,2^ biophysics,^3,4^ and environmental science.^5^ However, the ability to measure the behavior of particles in microfluidic cross-slot junctions is critically dependent on the precision, reproducibility, and cross-section uniformity of the channel geometries.^6^ Microfluidic channels with highly consistent dimensions and non-deformable walls under high pressures are essential for reliable experimental outcomes.^7^

However, empirical optimization of microfluidic cross-slot junction design can be infeasible, as achieving the necessary precision in rigid microfluidics presents significant challenges.^8^ For a single design, sufficient precision in all dimensions can be achieved with injection molding. However, the high cost of manufacturing injection molding tooling for a diverse set of designs can be prohibitive, even when extremely high precision is not required.^9^ Furthermore, the fabrication of multiple multi-layer photolithography masters with layer thickness controlled within 100 nm and subsequent translation of those features to nickel shims for injection molding is exceedingly challenging.^10^ Other prototyping methods for rigid microfluidics, such as casting in epoxy, direct 3D printing, and glass or glass-silicon chips each have their own limitations.^8^

Computational fluid dynamics (CFD) provides a powerful tool for optimizing channel geometries, focusing on metrics such as the frequency and amplitude of the oscillatory trajectories of particles exiting the cross-slot junction.^6,11^ Our own simulations have identified stenosis width and outlet width as critical parameters influencing particle trajectories. However, experimental validation of these simulated results has so far remained a challenge due to the sensitivity of particle behavior to the channel height, which thus must be tightly controlled while varying the geometry in width (x) and length (y) dimensions (Figure 1e).^6^

**Figure 1:**
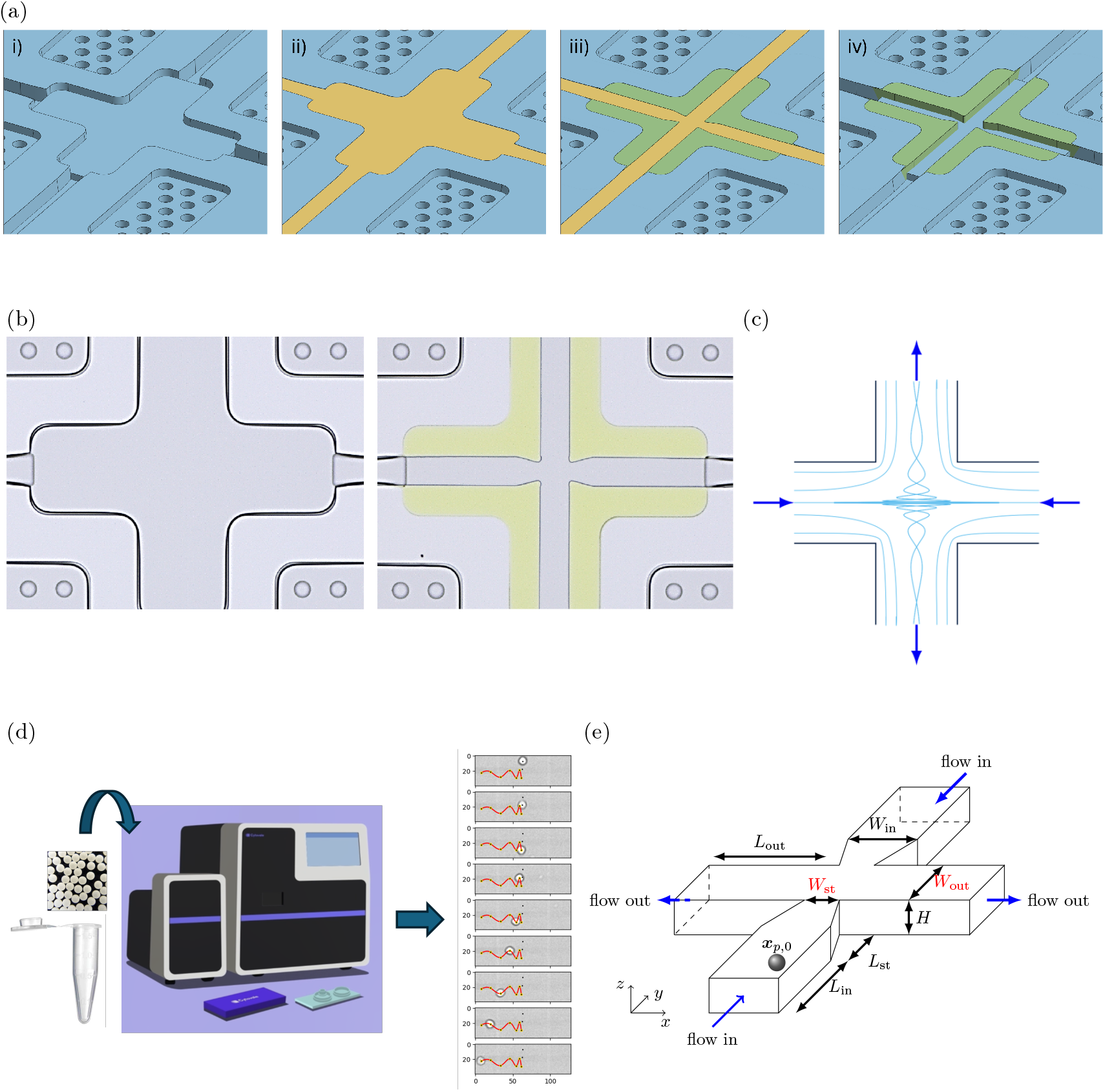
Overview of fabrication, simulation, and experimental methods. (a) Illustration of 2PP 3D printing process. i) Sealed microfluidic chips are produced with a cavity where the feature of interest will be printed (note: cover chip transparent in illustration); ii) Liquid photopolymer (yellow) is introduced into the sealed channel; iii) Regions outside the desired channel (green) are polymerized using 2PP 3D printing; iv) Unpolymerized photopolymer is flushed from the channel. (b) Optical microscopy images of injection molded and bonded ‘junction-less’ microfluidic chip before and after 2PP 3D printing of junction geometry. 2D projection of streamlines in vortex that forms in cross-slot junctions under specific flow conditions. Workflow of experimental validation of CFD model. A polystyrene bead solution is introduced into a cartridge containing a prototype microfluidic chip and processed on the Cytovale System, which captures and analyzes high-speed video to quantify particle trajectories through a microfluidic cross-slot. (e) Schematic of CFD simulation setup. Variable dimensions, stenosis width (*W*_*st*_) and outlet channel width (*W*_*out*_), are highlighted in red (Section 4.6). The initial position of the bead, *x*_*p*,0_, is represented by the gray sphere.

In this study, we provide the first experimental validation of CFD-predicted behaviors of micron-scale particles within microfluidic cross-slot channels. This advancement was made possible by a novel manufacturing approach enabling the production of microfluidic chips with exceptionally precise channel geometries. Injection molding from a single tool, combined with a tightly controlled solvent-assisted bonding process, achieved a channel height standard deviation of only 40 *nm* across ten cross-slot designs. Additionally, building on previous work demonstrating two-photon polymerization (2PP) 3D printing through a cyclic olefin polymer (COP) film,^12,13^ we applied this method to print through a 1 mm-thick COP cover slide to integrate sub-micron-resolution 2.5D cross-slot geometries within molded cavities in bonded chips (Figures 1a and 1b). The thick cover slide enabled the use of the high pressures necessary to achieve vortex formation in the cross-slot junction (Figure 1c) without channel deformation. Using the Cytovale System, a commercial platform developed to detect disease based on measurements of cell deformability,^1,2^ we validated the CFD model by analyzing the trajectories of polystyrene microspheres as a model for rigid particles (Figure 1d), demonstrating the effect of varying stenosis and outlet channel widths on particle behavior (Figure 1e).

In this work, we describe the method used to manufacture prototype chips with cross-slot geometries, a thorough metrological characterization of the dimensions of critical features on the prototype chips, the results of simulations of fluid and particles in corresponding geometries, and an experimental validation of these simulations. This work establishes a critical framework for the precise experimental investigation of particle behavior in microfluidic systems and highlights the synergy between advanced manufacturing techniques and computational modeling for optimizing microfluidic designs.

## 2 Results

### 2.1 Choice of Variables for Optimization

The performance of the cross-slot is influenced by several geometric parameters, but this study focuses on those most likely to affect the strength and structure of the vortex which forms in the outlet channels.

Two geometric features are varied: the outlet channel aspect ratio, *α*_out_ = *W*_out_*/H*, and the stenosis aspect ratio, *α*_st_ = *W*_st_*/H*. The outlet aspect ratio is known to influence the vortex dynamics,^14, 15^ while the stenosis aspect ratio controls the velocity of the impinging jets that drive the vortex. The height of the device, *H*, is fixed by fabrication constraints and is not varied.

The design space explored in the experiments is limited to five discrete levels for each parameter, corresponding to ±10% and ±20% variations around a baseline geometry.

### 2.2 CFD Simulations

#### 2.2.1 Fluid Vortex Characterization

Figure 2 shows slices of the vorticity magnitude scaled by the fluid advection time (|*ω*|*t*_adv_) in the *x*-*y* plane at *z* = *H/*2 for a sample of the simulations. The top row shows variation with the outlet aspect ratio, and the bottom row shows variation with the stenosis aspect ratio. Figure 2 uses the same color scale for all slices, allowing us to directly compare the vorticity magnitude between the different simulations.

**Figure 2:**
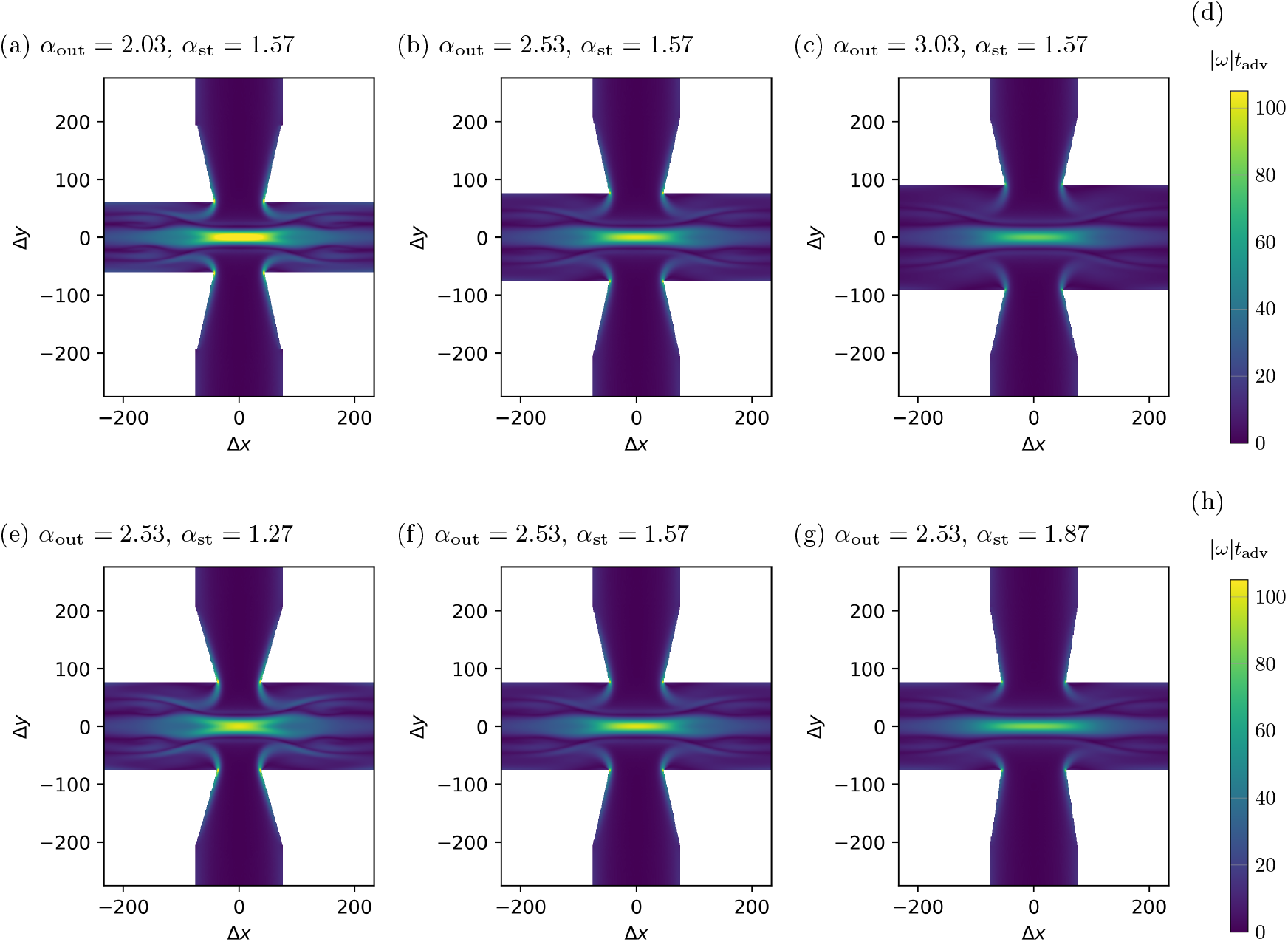
Simulation cross-sections showing the variation of the vorticity magnitude scaled by the fluid advection time (|*ω*| *t*_adv_) in the *xy*-plane at *z* = *H/*2. All use the same color scale. The top row shows variation with the outlet aspect ratio, and the bottom row shows variation with the stenosis aspect ratio. Panel (d) shows the color bar for the vorticity magnitude, repeated in panel (h) for clarity.

Looking at the top row, from panel (a) to panel (c), we see that as the outlet aspect ratio is increased, the vorticity magnitude is reduced. This behavior is likely due to the reduced fluid velocity in the outlet channel, which may reduce the intensity of the fluid vortex. Looking at the bottom row, from panel (d) to panel (f), we see that as the stenosis aspect ratio is increased, the vorticity magnitude also decreases. This trend is likely due to the the increased stenosis aspect ratio leading to a lower fluid velocity at the junction entrance, which in turn imparts less kinetic energy to the vortex.

These observations will be used to inform the interpretation of the bead behavior in the following sections.

#### 2.2.2 Simulated Trajectory Metrics

To quantify the differences in the trajectories as a function of the outlet width and stenosis width, we consider three metrics which have previously been used to characterize bead trajectories in cross-slot junctions,^6^ namely the residence time, the average angular velocity of the bead centroid around the outlet axis, and the number of turning points in the trajectory. Figure 3 shows the variation of these metrics with changes in outlet aspect ratio and stenosis aspect ratio for our simulations. Also included is a linear fit to the data, which is shown as a dashed line. For a visual representation of the trajectories themselves, see Figure S2.

**Figure 3:**
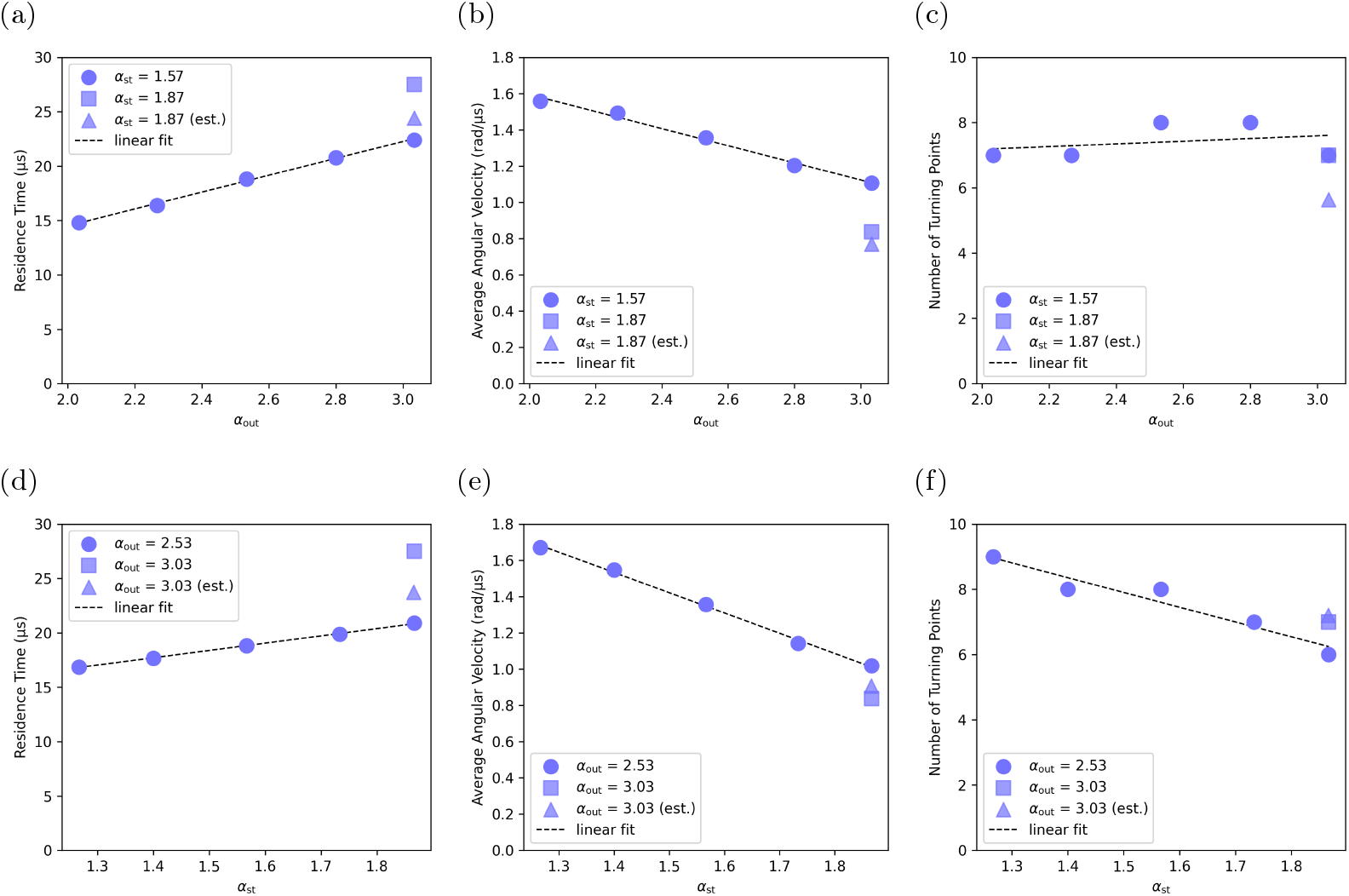
Metrics derived from simulated particle trajectories as a function of geometry. The top row (a-c) illustrates changes with outlet aspect ratio (*α*_out_), while the bottom row (d-f) shows variations with stenosis width (*α*_st_). Metrics displayed from left to right are: residence time, angular velocity (rotations), and number of turning points. Circular markers indicate single-parameter variations (top row: varying *α*_out_ with *α*_st_ fixed at 1.57; bottom row: varying *α*_st_ with *α*_out_ fixed at 2.53), with linear fits shown as dashed lines. The square marker shows the case where both parameters are maximized (*α*_out_ = 3.03, *α*_st_ = 1.87). The triangular markers show the estimated values for the metrics where both parameters are maximized, calculated from the linear fits of the metrics at fixed stenosis and outlet aspect ratios.

The residence time is defined as the time which a bead’s centroid spends in the junction region (Section 4.6.1). It is intuitive to expect the residence time to increase with increasing outlet aspect ratio, as a higher aspect ratio, i.e., wider channel, will result in lower average fluid velocity in the outlet channel. The behavior of the residence time with changes in stenosis aspect ratio is less intuitive, but is likely due to a narrower stenosis causing beads to enter the junction closer to the lateral center of the junction, which has previously been shown to result in a higher residence time^6^ due to the competing effects of drag from each outlet direction being balanced.

The average angular velocity is the average angular velocity of the bead’s centroid around the outlet axis while it is in the junction region. Similar to the residence time, the average angular velocity is linearly dependent on the outlet and stenosis aspect ratios. However, unlike the residence time, the average angular velocity is negatively correlated with the outlet aspect ratio and the stenosis aspect ratio. We hypothesize that the negative correlation of the average bead angular velocity with the outlet aspect ratio is due to the decreased average fluid velocity in the outlet channel leading to a reduced intensity of the fluid vortex. The negative correlation of the average angular velocity with the stenosis aspect ratio is likely due to a larger stenosis width resulting in a lower fluid velocity at the inlet to the junction, which is also likely to reduce the intensity of the fluid vortex.

The number of turning points in the trajectory is the number of turning points (defined in Section 4.5.1) which a bead completes while in the junction region. The number of turning points is not as clearly linear with respect to channel aspect ratios as the residence time or average angular velocity, but still shows an approximate linear relationship with both the outlet and stenosis aspect ratios, which is likely due to the integer constraint on the number of turning points. The number of turning points has no strong correlation with the outlet aspect ratio and a negative correlation with the stenosis aspect ratio. We can use the model given by,^6^ which states that the number of turning points is linearly correlated with both the residence time and the average angular velocity, to explain these trends. Increasing the outlet aspect ratio results in an increased residence time and decreased average angular velocity, which will cancel one another out, resulting in a smaller dependence of the number of turning points on the outlet aspect ratio. Increasing the stenosis width results in a smaller increase in the residence time and a larger decrease in the average angular velocity, which will result in a negative correlation between the number of turning points and the stenosis aspect ratio.

The case where both the inlet and outlet aspect ratios are increased is shown as a square in each panel of Figure 3. To assess whether the effects of changing both parameters are additive, we also show an estimated value based on the linear fits of the metrics at fixed stenosis and outlet aspect ratios, which are shown as a triangle marker on each panel. If the effects of changing both parameters were additive, then the estimated (triangle) values should be equal to the measured (square) values. Clearly, this is not the case, which suggests that the effect of changing both parameters is not purely additive.

### 2.3 Microfluidic Chip Characterization

The utility of 2PP 3D printing as a prototyping method to validate CFD models of microfluidic chips is dependent on its ability to form channels that meet the assumptions of the CFD models. We therefore employed multiple analysis techniques to robustly evaluate the precision, accuracy, uniformity, and limitations of 2PP 3D printing inside sealed, injection-molded microfluidic chips. We used collimated white light interferometry to evaluate the height of the channels, optical microscopy and SEM to evaluate the widths of the printed channels and features, optical microscopy to evaluate print alignment, and SEM to evaluate the shape of the printed channels.

#### 2.3.1 Channel Height

One of the key motivations to prototype junction geometries by printing 2.5D channels within injection molded chips was to remove differences in channel height as a variable. The variability in channel height that would result from fabricating a unique nickel insert for each geometry would result in an inability to attribute observations of differences in functional performance metrics to differences in target feature dimensions, as these differences could also be attributed to differences in channel height.

The height of both the junction and the inlet and outlet channels were measured on one chip of each geometry (*n* = 10 total; Figure 4a-c). For the junction measurements, height was averaged over the entire area of the shallow channel region, or junction layer; as such, the measurement area differs between junction geometries (wider channel designs have a larger measurement area than narrower channel designs). Because the channel height is not perfectly uniform across the junction (Figure 4a), this resulted in slightly elevated variability in the average height of the junction region relative to the variability in the deeper inlet and outlet channel regions.

**Figure 4:**
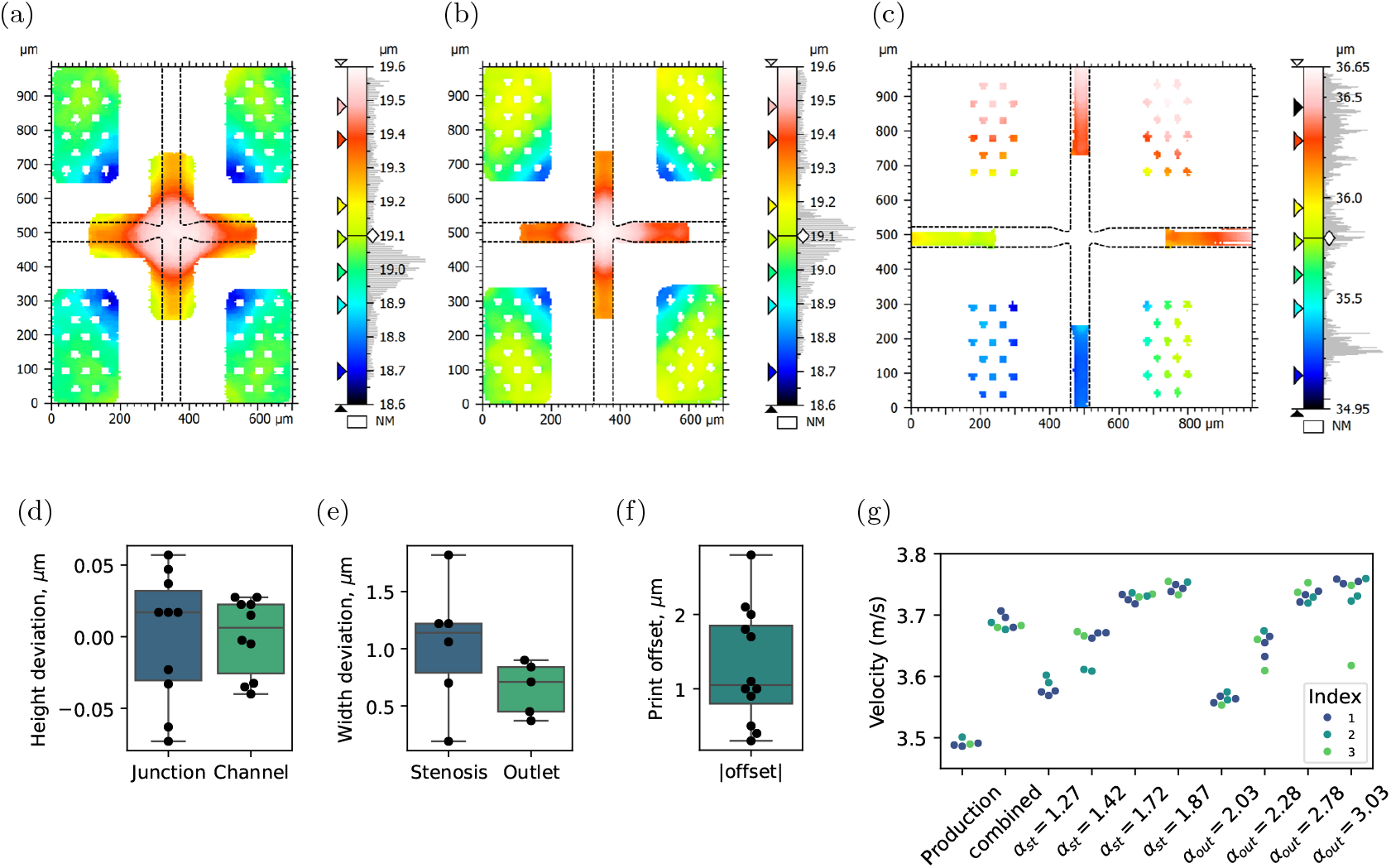
Characterization of prototype microfluidic chips. Panels (a–c) show example channel height measurements using collimated white light interferometry. Panels (d–g) present quantitative precision and accuracy characterization across multiple chips. (a) Junction layer measured after bonding but before 2PP 3D printing. (b) Junction layer measured after 2PP 3D printing. (Note: (a) and (b) are from different chips, but both were molded and bonded in the same batch.) (c) Feed/outlet layer. Slight height differences between the four channels result from spin-coating a thick photoresist layer. (d) Deviation in channel height from the mean for *n* = 10 prototype chips, shown separately for the junction region (blue) and the feed/outlet region (green). (e) Deviation from target dimensions in stenosis width (blue, *n* = 6) and outlet width (green, *n* = 5). (f) Offset between the printed cross-slot center and the center of the molded cavity. (g) Bead velocity in different prototype designs. Higher velocities correlate with higher junction aspect ratios and thus lower fluidic resistance. Velocity variability reflects structural repeatability. Commercial ‘Production’ chips produced via injection molding are shown for comparison. Each color indicates a unique chip.

For the junction region, the channel height standard deviation was 0.04 *µ*m, just 0.23% of the mean channel height. For all four inlet and outlet regions, channel height standard deviation is *<*0.1% of the channel height (Figure 4d).

#### 2.3.2 Print Accuracy and Precision

The dimensions of 2PP 3D-printed structures within the injection molded chips were accurate and highly precise. We evaluated print accuracy by comparing the measured width of printed features with the target width. We evaluated print precision by analyzing the variability in measurements of common features in subsets of chips (i.e., measurements of outlet width on chips with varying stenosis width and measurements of stenosis width on chips with varying outlet width).

Outlet width was measured on five chips with a common target channel width (Figure 4e). The standard deviation of the measured outlet channel width on these chips was 0.23 *µ*m, just 0.5% of the mean outlet width. The mean difference between measured and target width was 0.65 *µ*m which represents a mean error of 1.3% of the target outlet width. The range of error in outlet channel width was [0.37 *µ*m, 0.90 *µ*m].

Stenosis width was measured on six chips with a common target channel width (Figure 4e). The standard deviation of the measured stenosis width on these chips was 0.55 *µ*m, just 1.8% of the mean stenosis width. The mean difference between measured and target width was 1.04 *µ*m which represents a mean error of 3.5% of the target stenosis width. The range of error in stenosis width was [0.19 *µ*m, 1.82 *µ*m].

We observed no correlation (*r*^2^ = 0.146) between the magnitude of the difference between measured and target channel widths and the measured channel width, indicating that the absolute error is independent of the size of the feature.

#### 2.3.3 Print Alignment

From optical images, we compared the position of the center of the junction calculated using the boundary between the printed structures and injection molded chip and the position of the center of the junction calculated using the edges of the printed channels (Figure S1). The mean and standard deviation of the magnitude of the vector offset of the printed structures relative to the injection molded chip was 1.30 ± 0.74 *µ*m (Figure 4f).

#### 2.3.4 Channel Cross-Sectional Area

The CFD model assumes perfectly straight channel walls. To assess how closely the physical chips meet this assumption, SEM images of channel cross sections were analyzed. One chip of each geometry variation (and two chips of the geometry variation in which both outlet width and stenosis width were varied) was sectioned along the channel axis perpendicular to the feature with a unique width. Across all designs, a slight curvature in the channel walls was observed (Figure 5). To quantify the non-ideality of the channel cross-section shape, the area expected for a perfectly rectangular channel (calculated by multiplying the height and width extents of the channel cross-section) was compared to the cross-sectional area measured directly (via particle analysis).

**Figure 5:**
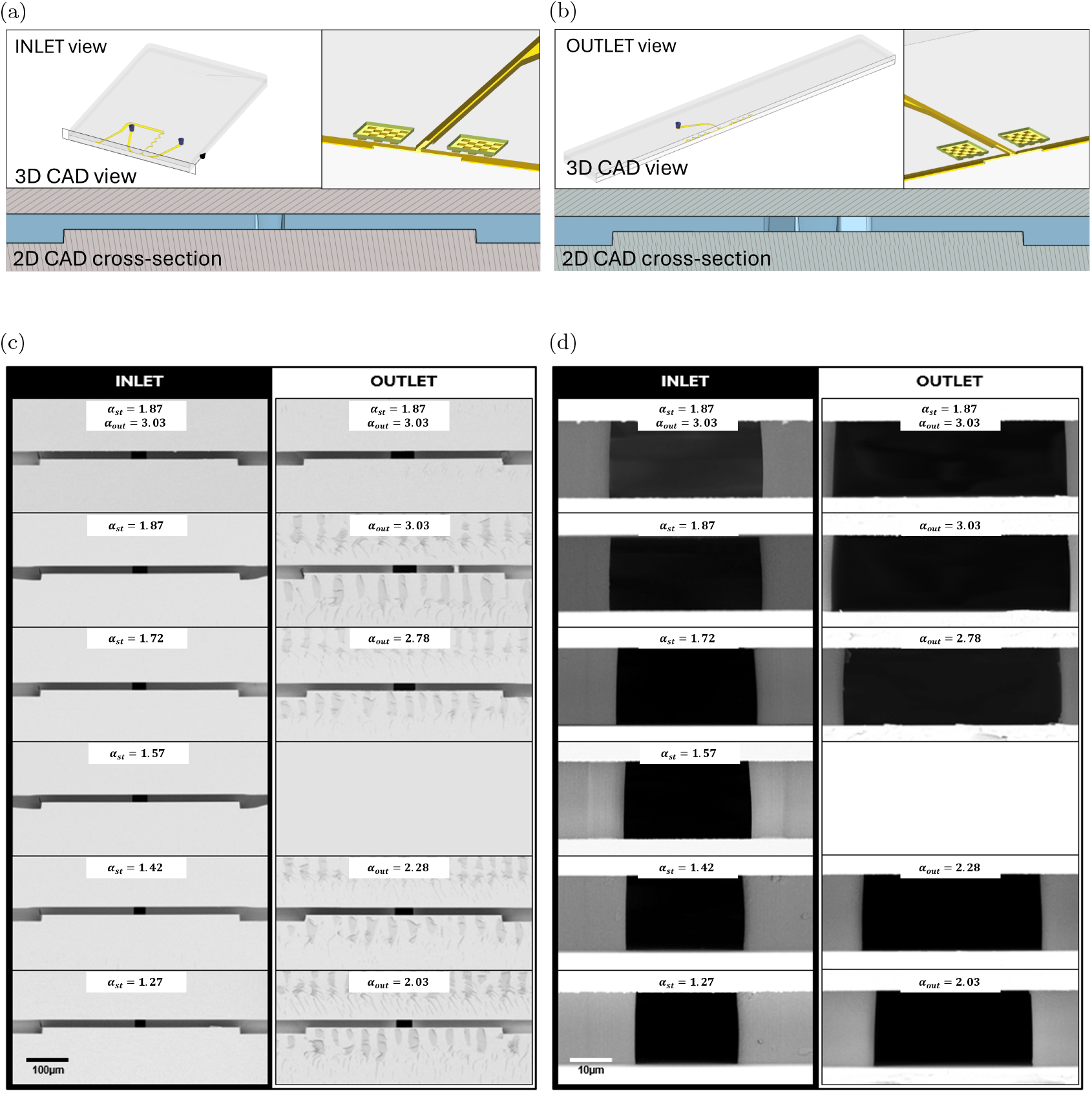
SEM imaging of prototype chips. Chips were sectioned along either the outlet channel centerline and imaged from the junction looking into the inlet channel (a, INLET) or sectioned along the inlet channel centerline and imaged from the junction looking into the outlet channel (b, OUTLET). SEM cross-sections of chips printed with each printed geometry at 800x magnification (c) and 1800x magnification (d).

The mean and standard deviation of the difference between the product of the channel cross section extents and the measured cross-sectional area were 33.3 ± 9.8 *µ*m^2^. This represents 4.2 ± 1.2 % of the cross-sectional area, meaning that the measured area is, on average, just 4.2% less than that which would be expected for perfectly rectangular channels. Additionally, the difference between these values was not correlated with the cross-sectional area (*r*^2^ = 0.081).

#### 2.3.5 Bead Velocity as Proxy for Geometry Consistency Across Chips

The velocity of beads through the chip is primarily dependent on the fluidic resistance of the chip and varies predictably with higher velocities measured in chips with wider channels (Figure 4g). As the channel height of the prototype chips is very tightly controlled, the fluidic resistance depends primarily on variability in the printed structures. Velocity variability is thus representative of overall geometric consistency of the printed structures.

For each individual chip, we considered the velocity measurement from the first run on that chip (Figure 4g). Each velocity measurement indicates the mean of individual particle velocities for a run measured immediately prior to the start of that run; velocities ranged from 3.55 to 3.80 m/s. We then calculated the standard deviation of the velocities for runs on three chips of each design (except for ‘*α*_*st*_ = 2.03’ for which runs were successful on only two chips). The median velocity standard deviation for chips with printed designs was 0.014 m/s, while the average velocity standard deviation for ‘production’ chips (injection molded junctions) was 0.008 m/s. The latter reflects variability in velocity not attributable to variability in printing, such as variability in channel height or inlet pressure. Of note, however, is that the range of standard deviations for velocity on chips with printed designs was [0.004, 0.034], indicating that in some cases, print reproducibility across three chips was comparable to that of chips with an injection molded junction.

### 2.4 Experimental Validation of CFD Simulations

Using the method described in Section 4.5.1, we estimated the turning points from the experimental data. Figure 6 shows the experimental turning points compared with the corresponding bead trajectories. The experimental data and numerically predicted trajectories are in reasonable agreement, with the amplitude and timing of the turning points on average deviating by 25% and 16%, respectively.

**Figure 6:**
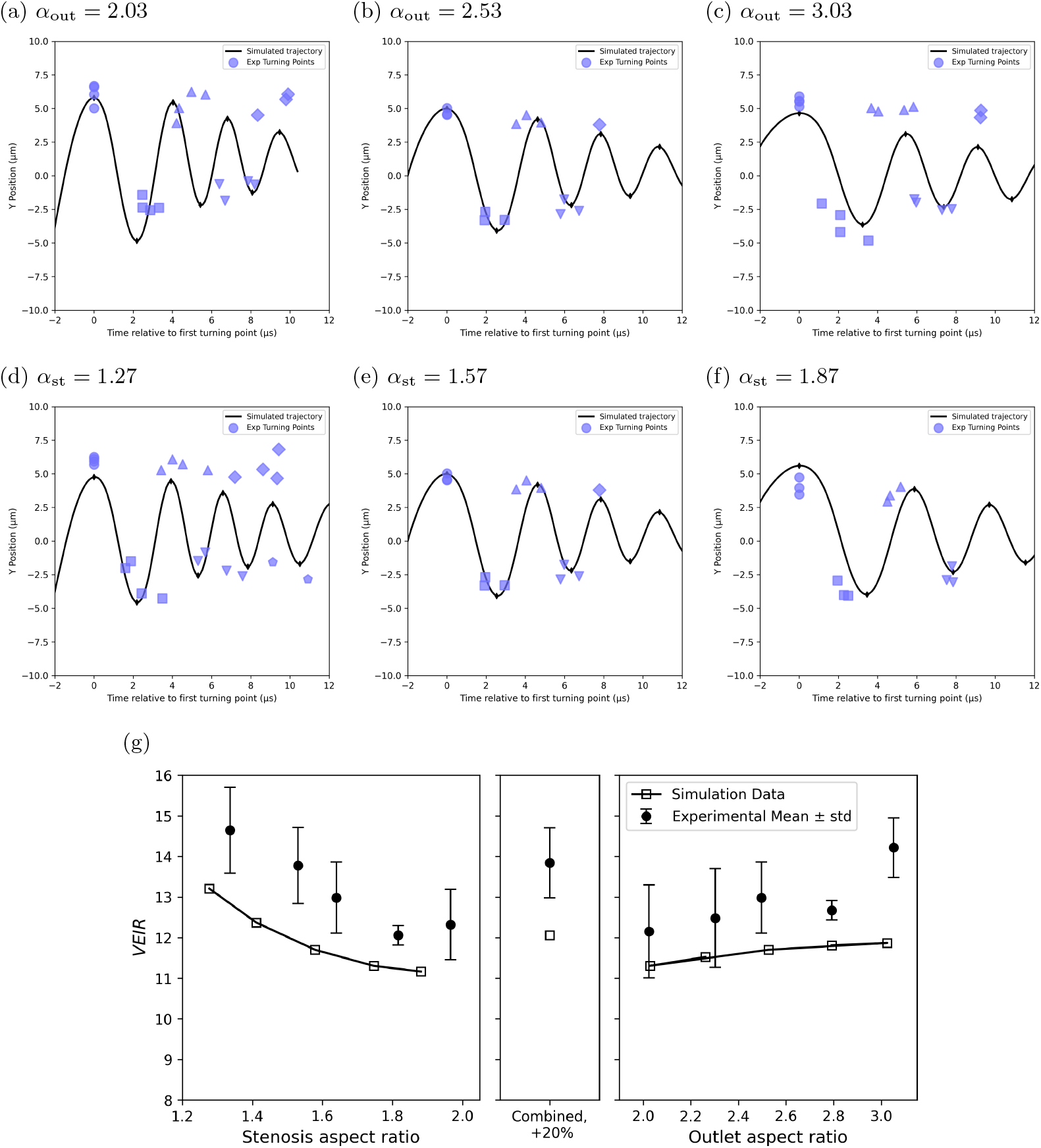
Comparison of experimental and simulated particle trajectories. The *y*-coordinate of turning points from experimental data (blue) and simulated trajectories (black) are plotted against time for various channel geometries. For subplots (a-c), outlet aspect ratio is fixed (*α*_out_ = 2.53), and for subplots (d-f), stenosis aspect ratio is fixed (*α*_st_ = 1.57). The time scale is given relative to the time of the first turning point. Different marker shapes are used for different turning point indices. Panel (g) shows simulated and experimental trends in VEIR, where with increasing stenosis aspect ratio, VEIR decreases (with the gradient decreasing with increasing stenosis aspect ratio), and with increasing aspect ratio, VEIR increases.

To demonstrate the utility of the simulations, we used a clinically-relevant metric, VEIR, that describes the amplitude of the particle trajectories (Figure 7c). Specifically, we compared VEIR predicted by simulations to VEIR measured from experimental data. Figure 6g shows that VEIR is predicted to decrease monotonically with stenosis aspect ratio, and increase monotonically with outlet aspect ratio. VEIR is more sensitive to the outlet aspect ratio than the stenosis aspect ratio, which is likely due to the larger magnitude of the outlet aspect ratio.

**Figure 7:**
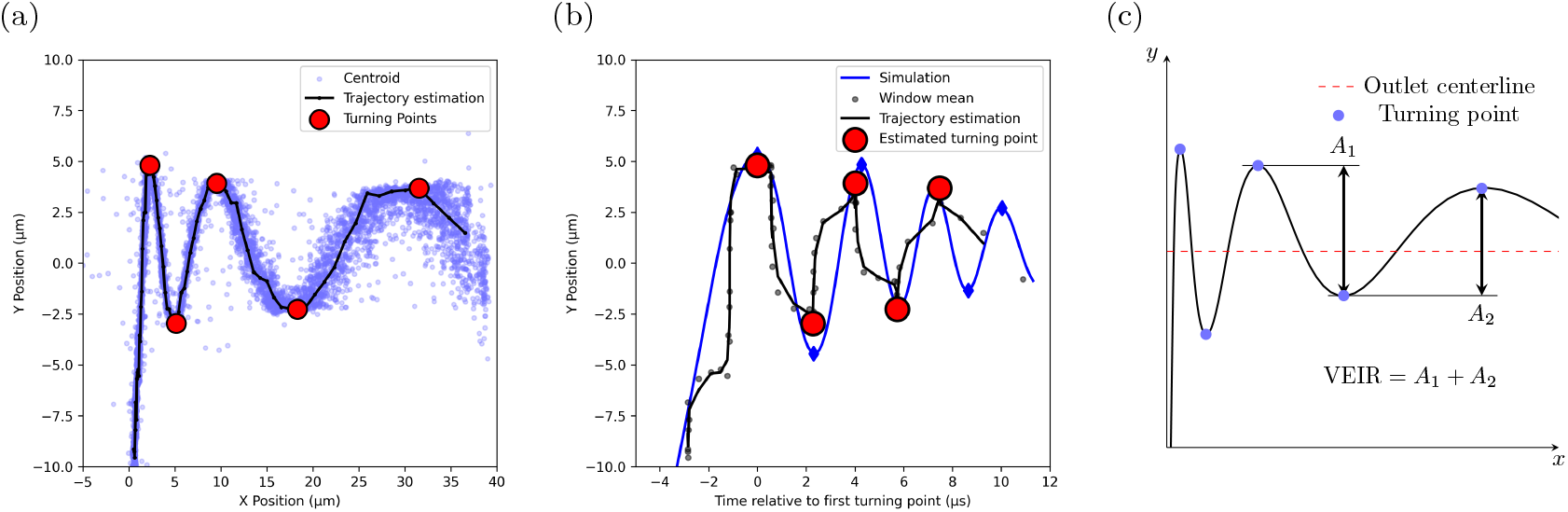
(a) Illustration of the method used to recover trajectories from point clouds of all measured bead centroids. (b) Illustration of the estimated trajectory from panel (a) plotted as lateral position vs. time. The points are the mean of the averaging window used to estimate the trajectory. (c) Illustrative definition of the viscoelastic inertial response (VEIR) metric and how it is calculated from a bead trajectory.

The experimental VEIR was consistently about 8% higher than the predicted values, but the trend of VEIR with respect to the aspect ratios is predicted well by the simulations. Error between the simulated and experimental VEIR values is shown in Figure S3.

## 3 Discussion

This work represents a novel application of highly-accurate 2PP 3D printing of structures inside of sealed, injection molded microfluidic channels. Pioneering work demonstrating printing inside COP channels used devices that were hot embossed,^12,13^ a process that is less reproducible and lower throughput than injection molding. Additionally, previous demonstrations used devices sealed with a thin COP foil, which would be expected to deform at the high pressures used in the Cytovale System. Printing with high accuracy through the relatively thick COP slide necessary for this application is inherently more challenging than either printing through a COP film or printing inside of PDMS channels bonded to a cover slip for several reasons: the requirement to use an objective with longer working distance to print through 1 mm-thick COP, reduced flatness of molded parts, lower effective resolution due to surface finish and potential bonding-induced internal stresses, and poor heat dissipation of COP resulting in localized heating of the resin.

We show that a high degree of precision and accuracy can be obtained even when 2PP 3D printing through a 1 mm-thick COP slide in sealed channels. Using multiple metrology techniques, we demonstrate that highly reproducible and accurate structures can be realized via this method. Sub-micron accuracy was achieved in channel width, the feature under investigation, and accuracy could be improved further were another batch of chips to be printed by compensating the printed design by the +0.65 *µ*m mean offset measured via optical microscopy in the outlet channel width.

As a functional demonstration of the reproducibility of printing, velocity measurements show variability approaching that of chips on which all features were injection molded, the gold standard for feature reproducibility in rigid microfluidic devices. This reproducibility is made possible by relying on established injection molding and solvent bonding processes to achieve channel height consistency across printed junction designs not possible via other methods. Even if standard SU-8 photolithography and PDMS soft lithography processes were used to prototype chips with different junction designs, *even if* all designs were patterned on the same wafer, achieving a standard deviation in channel height of just 40 *n*m on channels large enough to permit leukocyte transit would approach impossibility. We know of no other method with which 10 different 2.5D microfluidic channel designs could be prototyped with such consistent channel heights.

Using 2PP 3D printing to form critical structures has the additional benefit over injection molding of near-vertical sidewalls, as releasing parts from a mold with near-vertical sidewalls without introducing mold release defects is not always possible.^16^ It should be noted, however, that the ideality of the channel cross-section shape for 3D printed structures may decrease as channel height increases.

The ability of a simulation to predict real-world performance is only possible if the realized geometry closely matches the simulated geometry. The agreement of VEIR predicted by simulations and measured experimentally on chips with printed features (Figure 6g) demonstrates that 2PP 3D printing inside sealed thermoplastic chips is sufficiently reproducible to enable the experimental validation of a lattice-Boltzmann CFD model of rigid particle trajectories in a microfluidic cross slot. This model accurately predicts particle trajectories as the cross-slot geometry is varied, providing a powerful tool for the optimization of cross-slot geometries to achieve specific particle trajectories.

The offset observed between simulated and measured VEIR has a few possible explanations and illustrates limitations of this technique (and, more broadly, experimental validation of CFD simulations in microfluidics). First, simulations assumed a uniform channel height throughout the junction. As illustrated in Figure 4, channel height, and therefore channel aspect ratio, varies radially within an approximately 300 nm range with distance from the junction center. Second, simulations were run assuming a channel aspect ratio equal to the target, but the measured stenosis aspect ratio differed by up to 4.0% and the measured outlet channel aspect ratio differed by up to 2.0%. Third, while the channel sidewalls were nearly vertical, they were not perfectly vertical, and the simulation does not take into account the effect of non-vertical channel sidewalls. Fourth, simulated geometries lack radii on the channel wall corners in the junction, the effect of which is difficult to predict. And finally, simulations assumed a fixed volumetric flow rate, but experimental data was collected on a pressure-driven system with inlet pressure held constant. As illustrated in Figure 4g, velocity (as measured upstream of the junction in a channel with fixed dimensions across all designs) and therefore volumetric flow rate are design-dependent.

Differences in VEIR were significant (p *<* 0.01, unpaired t-test) between stenosis aspect ratios of 1.97 and 1.53, 1.97 and 1.34, 1.82 and 1.53, and 1.82 and 1.34, and between outlet aspect ratios of 3.05 and 2.30 and 3.05 and 2.02.

Microfluidic devices for applications where high pressures (*>*30 psi) are required, such as high-throughput droplet generation, have historically been difficult to prototype. This is especially true if the device has additional requirements, such as precise channel geometry and optical clarity or low autofluorescence. Methods for prototyping these devices have significant drawbacks: cost and turn-around time of tooling for thermoplastic devices, cost and throughput for glass or glass-silicon chips, resolution and optical clarity for traditional 3D printing (e.g., SLA), and autofluoresence, throughput, and solvent compatibility for epoxy resins.

2PP 3D printing inside sealed thermoplastic microfluidic channels is a powerful tool for cost-effective rapid prototyping, especially when rigid channel walls and high-precision geometries are necessary. While the injection-molded and solvent-bonded chip used in this work is a custom design, it is also possible to print structures in commercially-available thermoplastic chips, making this approach exceptionally accessible and obviating the need for capital investments in micro-manufacturing equipment for prototyping. The work presented here thus represents an exciting step forward for the experimental validation of CFD models of particle behavior in microfluidics when extremely high precision and accuracy are needed.

## 4 Materials and Methods

### 4.1 Fabrication of Injection Molded Microfluidic Chips

#### 4.1.1 Mastering

Inserts containing microfluidic structures for later use in micro injection molding were mastered using Ultra Violet Lithographie, Galvanoformung, Abformung (UV-LIGA), a widely used industrial process. UV-LIGA has previously been described in detail.^17,18^

In short, a negative photo-resist layer was spin-coated on a silicon wafer and the included solvent vaporized in a pre-bake step. A mask with the pre-defined structure was then applied to the photoresist surface and exposed areas of photoresist were cross-linked with UV exposure. Uncured photoresist was removed from the wafer using photoresist developer to obtain the desired 3D structure of the channels. This complete process was performed twice in order to generate the final 2-layer structure on the silicon wafer.

To form the nickel molds, a thin nickel seed layer was first sputter coated on the surface, then a thicker nickel layer was electroplated onto the remaining structures. Next, the wafer was separated from the nickel and remaining photoresist was chemically removed. Finally, the nickel template was polished and cut into the final tool shape.

#### 4.1.2 Injection Molding

The finished microfluidic chip consists of two molded components, the ‘fluidic chip’ (containing microfluidic structures) and the ‘cover chip’ (containing inlet and outlet holes). Injection molding of each component was performed on an Engel e-Motion molding machine (Engel Austria GmbH, Schwertberg, Austria) with a cyclic olefin polymer (COP), a polymer class widely used for microfluidic diagnostic devices.^19^ Cylinder temperature was set to 190-260^*°*^C, mold temperature to 50-95^*°*^C with a screw speed of 20-60 rpm and an injection speed of 30-100 cm^3^/s. Injection and holding pressure were set to 30-60 MPa and the back pressure to 4-10 MPa.

#### 4.1.3 Solvent Bonding

Fabrication of the final microfluidic chip was done by solvent activated bonding of the fluidic and cover chips.^20, 21^ In short, the bonding surfaces of the chips were activated with a solvent, followed by a joining process in a bonding press.

### 4.2 Integration of Micro-Scale Geometries into Microfluidic Chips Using 2PP Technology

The methods employed by NanoVoxel GmbH describe the integration of micro-scale geometries into microfluidic chips through 2PP technology and subsequent removal of unpolymerized resin.

The micro-scale geometry was initially adjusted in a CAD file (OnShape) to account for the refractive index of the COP chip and the resin. The design was uniformly scaled to achieve the desired final dimensions post-printing. In the 3D printing software Think3D (UpNano, Vienna, Austria), a layer height of 1 µm, laser scanning speed of 300 mm/s, hatching distance of 0.5 µm, and laser power of 40 mW were selected. A 10x objective lens (Olympus, Tokyo, Japan) was utilized in the 2PP printer.

Prior to resin perfusion, each microfluidic channel was cleaned by flushing with isopropanol (IPA, Sigma-Aldrich). UpPhoto (UpNano, Vienna, Austria), a photosensitive resin, was then injected until the junction was completely filled. This resin fully cures during the 2PP process and does not require additional thermal or optical post-treatments.

The resin-filled chip was placed on the auto-leveling stage of the NanoOne (UpNano, Vienna, Austria). Using the Think3D software, the junction region was aligned in the x-y plane with the channel walls serving as calibration edges, minimizing translational and rotational misalignments. Three points within the junction were measured to detect the interface between resin and COP, allowing the stage’s leveling motors to ensure a uniform Z position. This alignment ensured proper bonding of printed structures to both the top and bottom surfaces of the channel.

The main printing process was initiated from the channel bottom upward, employing the specified laser power, scanning speed, and layer settings. Polymerization progress was monitored in real time using the printer’s imaging system to ensure stable and accurate fabrication.

After the printing process, unpolymerized resin was removed by flushing the channels multiple times with IPA. Residual solvent was subsequently expelled using filtered air under gentle pressure, leaving behind fully cured 2PP-printed microstructures.

### 4.3 Metrology

#### 4.3.1 Channel Height Metrology

Channel height data was acquired by interferometric measurement (FRT MicroProf, Fries Research & Technology GmbH, Bergisch Gladbach, Germany) of the fabricated chip. MountainsMap software (v.10.2.10719, 2024/05/07, Digital Surf) was used to calculate channel height from the raw data. Qualitative channel top and bottom geometry was acquired by confocal microscopy (Keyence VK-X250 CLSM, Mechelen, Belgium).

#### 4.3.2 Channel Width and Print Alignment Metrology

The channel width (inlet channel, outlet channel, stenosis) and print alignment (2PP vs. injection molded structures) were both measured using a high resolution Keyence digital microscope VHX-7000 (Keyence International, Belgium) in transmission illumination configuration, and using the E100 zoom objective lens at 500x magnification. The data analysis was performed using the VHX Remote Analysis Software (v.3.0.34.332, Keyence Corporation) and built-in measurement tools, with built-in edge detection algorithms. The x-y alignment accuracy between the 2PP and the injection-molded structures was measured by calculating the centerlines of the respective channel geometries, calculating the junction center points from the intersection points of the centerlines, and measuring the distance of the two junction center points (Figure S1).

#### 4.3.3 FE-SEM Imaging

For FE-SEM Imaging, sealed chips containing 2PP 3D-printed structures were cross-cut with a microtome (Type HM 355, MICROM GmbH, Walldorf, Germany). A conductive Au-layer was applied on the cross-cut surface with a sputter coating machine (SCD 050, BAL-TEC, Balzers, Liechtenstein) at 50 mA for 60 s in an argon plasma prior to SEM. Images were acquired at 2 kV EHT voltage, 10 mm working distance with a secondary electron detector (SE) at 800x and 1,800x magnification on a Zeiss Ultra 55 FE SEM (Carl Zeiss, Oberkochen, Germany) with SmartSEM (v.06.00.05, Carl Zeiss). MountainsMap software (v.10.2.10719, 2024/05/07, Digital Surf) was used to measure features on images.

Gold used in the sputtering was obtained from Materion, Mayfield Heights, OH, USA; argon was obtained from Air Liquide, Anif, Austria.

### 4.4 Consumables and Instrumentation

#### 4.4.1 Bead Solution

In place of a processed blood sample, a solution of 8 *µ*m-nominal diameter polystyrene beads (Duke Standards, Fremont, California, US) was prepared at a final concentration of 0.3% w/v in 0.01% Tween in PBS. This solution was prepared fresh for each day of testing and vortexed immediately before each run. Bead size was chosen to be close to that of white blood cells and match the ratio of particle and channel size used in simulations while minimizing the likelihood of clogs and disruption of the vortex in the junction.

#### 4.4.2 Consumables

Prototype chips, manufactured as described in Sections 4.1 and 4.2 were assembled into cartridges consisting of a pressure interface to the Cytovale System’s Cell Imaging Module, an inlet reservoir, an in-line mesh filter with a 10 *µ*m pore size, and an outlet reservoir. A hermetic seal was achieved with laser welding.

#### 4.4.3 Instrumentation

The Cytovale System contains three modules: the Sample Preparation Module (SPM), the Cell Imaging Module (CIM), and the Image Analysis Module (IAM). The SPM was bypassed for the experiments in which the data presented here was generated. The CIM consists primarily of a high-speed camera, illumination and optical system, and pneumatics system. The IAM consists of a workstation containing powerful GPUs and running custom software. The CIM and IAM were used to optically align the chip, apply 91 psi to the cartridge inlet, and record and store an 11.7 s video at 580,000 fps for each run.

#### 4.4.4 Experimental Procedure

Experiments were run on the Cytovale System according to the instructions for use for the IntelliSep test,^22^ with minor modifications. In place of a blood sample processed using the SPM, a bead solution was used,prepared as described in Section 4.4.1. Additionally, an engineering build of the Image Analysis Module software was used to enable aligning the center of the junction with the center of the field-of-view on chips with varying junction dimensions. Finally, instead of using the outputs of the Image Analysis Module directly, raw videos for each run were stored and reprocessed using an engineering data processing pipeline to obtain the inputs to the analysis presented in this paper.

For each run, 1 mL of bead solution was loaded into the cartridge inlet. For replicate runs on the same cartridge, after each run, a blunt-tip needle and syringe were used to remove excess sample from the cartridge inlet and outlet, with care taken to not damage the cartridge inlet filter.

For each chip design, three replicates were attempted on one cartridge and at least one replicate was attempted on each of two additional cartridges. As no difference in key functional performance metrics was observed between the first replicate and subsequent replicates on cartridges for which more than one replicate was successful, data from all replicates on a chip design were pooled, except when examining chip-to-chip variability within each design.

### 4.5 Data Analysis

#### 4.5.1 Estimation of Bead Trajectories and Turning Points

The raw data from each experiment comprises high-speed camera footage capturing beads as they transit the junction. For each frame where a bead is present, its outline is identified using a neural network, and this outline is processed to determine the bead’s centroid coordinates (*x*_*i*_, *y*_*i*_). The high frame rate yields approximately 10 centroid measurements per bead during a single transit.

Due to the steady-state nature of the flow field and the precise particle grouping achieved by upstream inertial focusing, a scatter plot overlaying all recorded bead centroids from an experiment reveals a well-defined helical trajectory shape (illustrated in Figure 7a). To estimate the locations of this trajectory’s turning points, we first transform the discrete point cloud of centroids into a continuous function *y* = *f* (*x*). This is achieved using a moving window approach (Figure 7b). For any given x-coordinate *x*, we consider all centroid points (*x*_*i*_, *y*_*i*_) that fall within a window of width *δ*, specifically where the *x*-coordinate *x*_*i*_ is between *x* − *δ/*2 and *x* + *δ/*2.

From the set of corresponding *y*-values *y*_*i*_ within this window, we compute a one-dimensional Gaussian Kernel Density Estimate (KDE). This KDE gives us a smooth probability distribution indicating where *y*-values are most likely to occur within that specific window. The function *f* (*x*) is then defined as the *y*-value where this calculated KDE reaches its maximum peak. In simpler terms, *f* (*x*) is the y-coordinate with the highest estimated density of points within the local *x*-window. By selecting a sufficiently small window width delta, this method provides a robust estimate of the underlying trajectory function *y* = *f* (*x*), from which the positions of the turning points can be readily calculated. The black line in Figure 7a represents the continuous trajectory.

Similarly, we can estimate the corresponding time for each point in the trajectory. If we define a time history which starts at the first detected image of a bead transiting the junction, we can average that time for every frame within a window used to estimate the trajectory, thus obtaining a time parameterization. Figure 7b illustrates this parameterization. Due to the discrete values of time corresponding to each frame, this averaging process is inherently noisy; taking a three point moving average reduces this noise.

In previous work, we presented evidence that a particle’s viscoelastic inertial response (VEIR; illustrative definition in Figure 7c) is predictive of sepsis in human patients.^2^ Here, we use VEIR as the primary metric for evaluating and comparing particle trajectories between experiments and simulations.

### 4.6 CFD Simulations

#### 4.6.1 Model Description

We use a simplified model of the cross-slot junction for simulations. A diagram of the simplified geometry is shown in Figure 1e. Importantly, we do not simulate the entire microfluidic chip, but only the junction and the channels immediately adjacent to the junction.

We assume a fully developed Poiseuille flow at the two inlet boundaries with constant average velocity, and a constant pressure open boundary at both outlet boundaries. Flow is Newtonian and incompressible. No-slip conditions are applied at the walls of the channel, and on the surface of the rigid bead.

We apply a constant inlet velocity for all studied geometries, upon the assumption that the junction region, as part of a much larger microfluidic device, has a minimal impact on the overall flow resistance.^23^

As we do not simulate the upstream bead focusing region of the chip, we assume that the bead is initially positioned at its vertical equilibrium position. We recover the vertical equilibrium *z*-position from simulations of a rigid bead in a periodic channel with the same aspect ratio and flow rate as the inlet channel of the junction. A detailed description of the setup of such simulations can be found in our previous works.^11,24,25^ The upstream inertial focusing of the bead should result in a tight grouping of beads at the lateral (*x* direction) center of the inlet channel. Perfect lateral focusing should result in a 50:50 distribution of beads leaving in the two outlet channels. However, we observe in experiments that nearly all beads exit the junction from the same outlet channel. Therefore, we assume the initial position of the bead has a slight lateral offset from the center of the inlet channel. For the sake of simplicity, we assume this offset is 0.1% of the channel width from the centerline.

We make some important assumptions in the geometry simplification, namely that the walls of the channel are perfectly straight, perpendicular (other than the angled stenosis section), and smooth. All filleting in corners is ignored, as is the draft angle of the vertical channel walls. Due to the numerical method being limited to solving the fluid in cubic voxels, the stenosis angle is approximated as a staircase, and all dimensions are approximated to the nearest voxel.

To make clear which areas are considered to be the inlet and outlet channels, and which is the junction, we define a ‘junction region’, which is a cuboid in the center of the junction with a height of *H*, extending a length of *W*_in_ in the outlet channel direction and *W*_out_ in the inlet channel direction.

#### 4.6.2 Numerical Methods

We solve the three-dimensional fluid flow using the lattice-Boltzmann method on a D3Q19 lattice. We select the BGK collision operator for simplicity, and the Guo forcing scheme to incorporate the external forces from the bead simulation. To satisfy the no-slip condition on the surface of the moving rigid bead, we use the multi-direct-forcing immersed boundary method with a three-point stencil. A detailed description of the model used in this work has previously been provided by Kechagidis *et al*.^6^

#### 4.6.3 Matching Simulations to Experiments

To make meaningful comparisons between experimental and simulated conditions, we make use of dimensionless quantities. This also allows us to directly compare simulations to experimental data, by using the same scaling.

We define several dimensionless aspect ratios using the channel height *H* as our reference length scale. The inlet channel aspect ratio is:

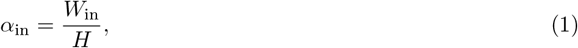

where *W*_in_ is the inlet channel width.

Similarly, the outlet channel aspect ratio is:

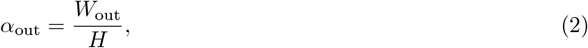

where *W*_out_ is the outlet channel width.

For the stenosis, we define its aspect ratio as the width at the narrowest part of the inlet nozzle relative to *H*:

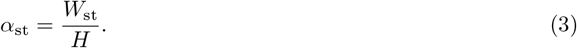

The dimensionless inlet nozzle length is obtained by:

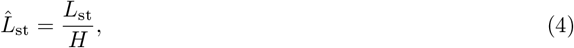

where *L*_st_ is the physical length of the inlet nozzle.

Finally, the particle confinement is defined as:

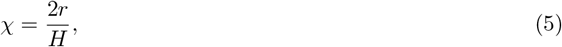

where *r* is the particle radius.

The Reynolds number of the inlet channel quantifies the influence of fluid inertia and is used to scale the fluid velocity. We use the inlet channel width as a characteristic length scale, as has been used elsewhere.^14^

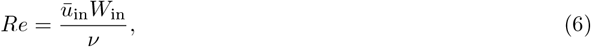

where 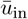 is the average velocity in the inlet channel and ν is the fluid kinematic viscosity.

To convert simulation time scales to experimental, we scale by the fluid advection time, *t*_adv_,

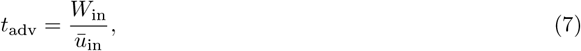

resulting in a non-dimensional time scale of

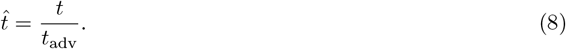

Each of the dimensionless quantities presented in this section is calculated for the experimental parameters, and is shown in Table 1. The simulation is then aligned to the experiment by choosing parameters which result in equal values for each dimensionless quantity, for a given lattice resolution fine enough for resolution independence.

**Table 1:** Comparison of key parameters used in simulation and experimental validation. Values shown represent the design parameters used for the geometry case where the outlet and stenosis aspect ratios are equal. ^†^Physical parameters, ^‡^dimensionless parameters, ^§^numerical parameter. All numerical parameters and simulation physical parameters are given in lattice units, relative to the lattice unit spacing Δ*x*, time step Δ*t* and unit lattice density. The simulation physical parameters are chosen such that they result in the dimensionless parameters being equal to the experimental values.

| Parameter | Simulation Value | Experimental Value |
| --- | --- | --- |
| Inlet channel aspect ratio ($\alpha_{\text{in}}$)<sup>‡</sup> | 2.53 | 2.53 |
| Outlet channel aspect ratio ($\alpha_{\text{out}}$)<sup>‡</sup> | 2.53 | 2.53 |
| Stenosis aspect ratio ($\alpha_{\text{st}}$)<sup>‡</sup> | 1.57 | 1.58 |
| Reduced inlet nozzle length ($\hat{L}_{\text{st}}$)<sup>‡</sup> | 2.2 | 2.27 |
| Particle confinement ($\chi$)<sup>‡</sup> | 0.42 | 0.42 |
| Reynolds number<sup>‡</sup> | 249 | 248 |
| Voxel size<sup>§</sup> | $\Delta x$ | 0.3167 $\mu\text{m}$ |
| Time step<sup>§</sup> | $\Delta t$ | 1.1 ns |
| Relaxation time ($\tau$)<sup>§</sup> | 0.53$\Delta t$ | — |

## Acknowledgments

Calum Mallorie received funding from Cytovale, and from the Engineering and Physical Sciences Research Council (EPSRC). Timm Krüger received funding from the European Research Council (ERC) under the European Union’s Horizon 2020 research and innovation program (803553).

## Conflict of interest statement

Authors CM, SdC, and TRC are affiliated with Cytovale and have an equity interest in the company whose system was used to evaluate the functional performance of prototype microfluidic chips. This system is covered by the following patents: 8,361,415; 8,935,098; 2,619,545; and 9,897,532.

Authors BS, MP, CH, and CL are affiliated with Stratec Consumables, which manufactured the assembled microfluidic chips.

Authors NM and DF are affiliated with NanoVoxel, which manufactured prototype geometries inside the assembled microfluidic chips. Author DF has an equity interest in NanoVoxel.

The other authors declare that they have no competing financial interests.

## Author Contributions

CM set up and ran all simulations, defined geometries to prototype, and analyzed experimental data. MP, BS, CH, and CL managed manufacturing of injection molded and bonded parts, conducted all metrology, and analyzed metrology data. NM and DF developed printing-in-chip method and printed all prototype junction geometries. SdC designed and ran polystyrene bead experiments. TK developed framework on which CFD simulations are based. TC ideated and managed the project and led writing of the manuscript.

## Additional Information

Supplementary Information is available for this paper.

