## Supplementary Information for "Optimization of Injection-Molded Thermoplastic Microfluidic Chip Design with Numerical Modeling and Two-Photon Polymerization 3D Printing"

### Measurement of Print Dimensions Using SEM vs. Optical

We measured feature width using two independent methods: optical microscopy of intact chips (Figure S1) and scanning electron microscopy (SEM) sectioned along the axis perpendicular to that being measured (see Methods). These two methods were highly correlated ($r^2$=0.998), though SEM measurements had an offset of +1.27 $\mu m$ compared to corresponding optical measurements. Optical measurements were used for reports of print accuracy and precision, as for SEM imaging, chips were cross-sectioned perpendicular to the variable feature, leading to n=1 measurements for each geometry.

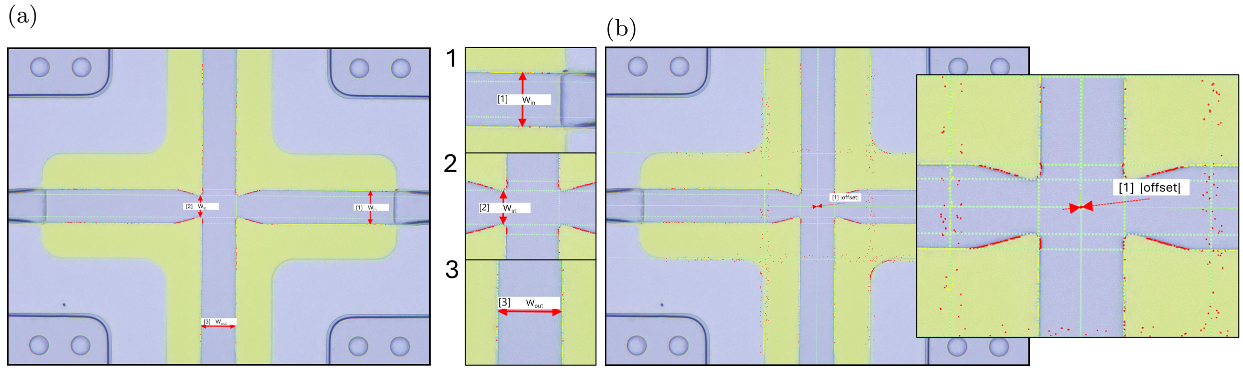


Figure S1: Measurement definitions for optical microscopy images. (a) 1. Inlet width ($W_{in}$) was measured as the distance between two lines parallel to one another and coincident with the edges of the printed inlet channel walls. 2. Stenosis width ($W_{st}$) was measured as the distance between two lines parallel to one another and the inlet channel walls and intersecting the most prominent point of the printed channel walls. 3. Outlet width ($W_{out}$) was measured as the distance between two lines parallel to one another and coincident with the printed outlet channel walls. (b) The offset between the printed and molded cross-slot center was measured as follows: the intersection of the printed channel centerlines (calculated from inlet and outlet width measurements in (a)) was used as the center position of the printed cross-slot; the center position of the molded structures was calculated in a similar fashion, but using the interface between the molded and printed structures as the inlet and outlet channel edges; the magnitude of the difference in position of these two center points is presented as the offset between printed and molded structures.

### Simulated Bead Trajectories

Figure S2 shows the path which a simulated bead takes through the center of the junction for a range of outlet width and stenosis width values. Each panel shows the path for a single outlet width value, with the other parameter varied. The dashed line shows the path for the case where both parameters set to the maximum value. From Figure S2, it is clear that changing both the outlet width and stenosis width results in a marked change in the path of the bead, but it is difficult to discern the individual effects of changing each parameter.

### Comparison of Simulated and Experimental VEIR

To evaluate how well our simulations capture experimentally observed trends in VEIR, we computed the relative error between simulated and experimental VEIR values for each chip design. Figure S3 shows the error as a function of aspect ratio, using the simulation value for alignment. Error bars represent the standard deviation across chips of the same design.

The relative error remains low and consistent across the range of aspect ratios, suggesting that the simulations accurately reflect how VEIR varies with channel geometry in the experimental data.

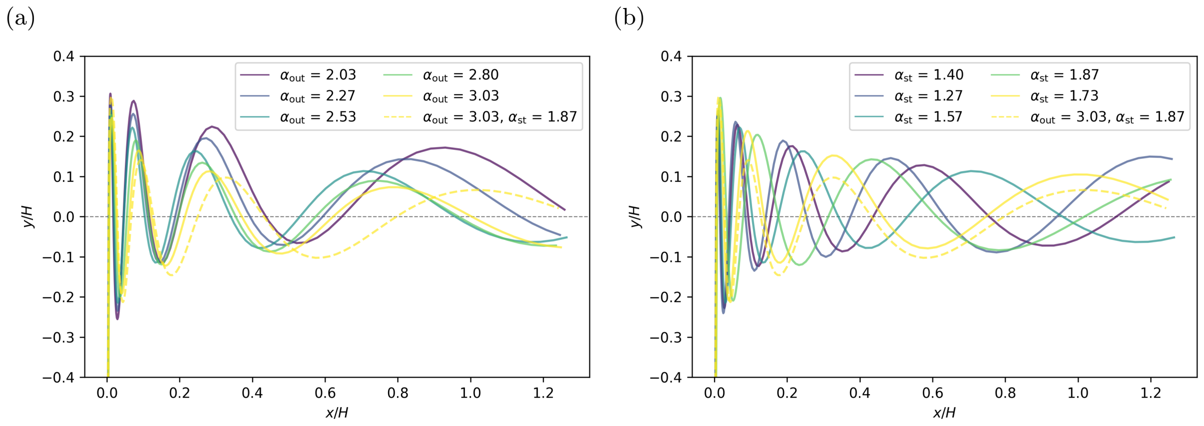


Figure S2: Simulation trajectories showing variation with geometry changes. Panel (a) depicts the variation of outlet aspect ratio changes, and panel (b) the stenosis aspect ratio. On each panel, we show the design with increased stenosis *and* outlet aspect ratios with a dashed line.

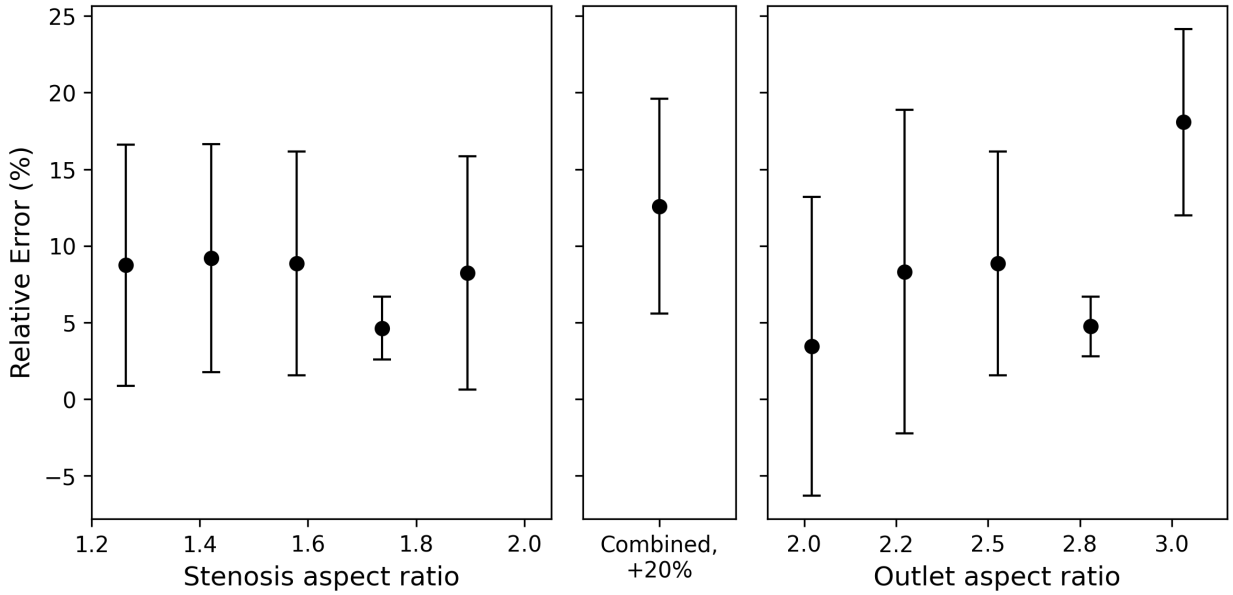


Figure S3: Relative error between simulated and experimental VEIR values for each chip design. The error is computed relative to the experimental value, with the aspect ratio corresponding to the simulation input. Each point represents a design, and error bars show the standard deviation across replicate chips. The absence of a systematic trend across aspect ratios indicates strong agreement between simulated predictions and experimental results.
